# Rethinking Alzheimer’s: Harnessing Cannabidiol to Modulate IDO and cGAS Pathways for Neuroinflammation Control

**DOI:** 10.1101/2025.03.06.641791

**Authors:** Sahar Emami Naeini, Bidhan Bhandari, Breanna Hill, Nayeli Perez-Morales, Hannah M Rogers, Hesam Khodadadi, Nancy Young, Lívia Maria Maciel, Jack C Yu, David C Hess, John C Morgan, Évila Lopes Salles, Lei P Wang, Babak Baban

**Author notes:** **Corresponding authors:** Babak Baban, Ph.D., -Department of Oral Biology, DCG., -DCG Center for Excellence in Research, Scholarship and Innovation (CERSI) Augusta University, Augusta, GA, USA.

## Abstract

Alzheimer’s disease has traditionally been associated with amyloid-β plaques, but growing evidence underscores the role of neuroinflammation in disease progression. The autoimmune hypothesis of Alzheimer’s disease suggests chronic inflammation and immune dysfunction contribute to neuronal damage, making modulation of immune responses a promising therapeutic strategy for the disease.

Cannabidiol, a phytocannabinoid with anti-inflammatory properties, may offer therapeutic potential. This study explores how cannabidiol influences the Indoleamine 2,3-dioxygenase (IDO) and cyclic GMP-AMP synthase (cGAS) pathway, a key regulator of neuroinflammation in Alzheimer’s disease.

Using the 5XFAD transgenic mouse model of Alzheimer’s disease, we administered cannabidiol via inhalation. We assessed immune markers, including Indoleamine 2,3-dioxygenase and cyclic GMP-AMP synthase, through flow cytometry, immunofluorescence staining, and gene expression analysis. Cytokine levels and neuroinflammatory responses were evaluated, and protein-protein interactions within the Indoleamine 2,3-dioxygenase/cyclic GMP-AMP synthase pathway were analyzed using the STRING database.

Cannabidiol treatment significantly reduced Indoleamine 2,3-dioxygenase and cyclic GMP-AMP synthase expression, correlating with lower levels of pro-inflammatory cytokines, including Tumor Necrosis Factor-alpha, Interleukin-1 beta, and Interferon-gamma. Bioinformatics analysis identified potential interactions between cannabidiol and immune targets such as Protein Kinase B (AKT1), Transient Receptor Potential Vanilloid 1, and G-protein coupled receptor 55, suggesting a multi-targeted therapeutic effect.

These findings support cannabidiol as a potential monotherapy or adjunctive treatment for Alzheimer’s disease, targeting neuroinflammatory pathways, particularly the Indoleamine 2,3-dioxygenase/cyclic GMP-AMP synthase axis. Further studies are needed to explore its full therapeutic potential.

## Introduction

Alzheimer’s Disease (AD) remains a major challenge for healthcare, with no effective treatment to halt or slow its progression (1-3). Current therapies, such as amyloid-targeting treatments and cognitive enhancers, only offer symptomatic relief, failing to address the root causes of neurodegeneration (3-4). This highlights a critical gap in our understanding.

The complexity of AD, driven by genetic, environmental, and immune factors, means no single theory can explain it fully. While amyloid-beta plaques and tau tangles are central to the traditional AD model, they do not account for the disease’s full scope (5-7). For instance, individuals with amyloid plaques may not exhibit cognitive decline, and some with severe neurodegeneration lack significant amyloid or tau pathology (8-9). This suggests the amyloid-tau framework is incomplete and may miss key aspects of AD’s true mechanisms.

Immune system dysfunction is considered a critical and obligatory step in the progression of AD. While amyloid-β (Aβ) accumulation is a key hallmark of AD pathology, recent evidence suggests that immune dysregulation can independently drive neurodegeneration. Specifically, the activation of immune responses, including microglial and astrocytic activation, has been shown to exacerbate neuroinflammation and contribute to neuronal damage. In the 5xFAD mouse model, while Aβ deposition occurs early in the disease, the immune system’s role in driving neurodegenerative processes extends beyond Aβ-related events. This aligns with the AD^2^ theory of AD, which posits that immune dysfunction, rather than being solely a consequence of Aβ accumulation, is an early and central driver of disease progression (10). Therefore, immune system dysfunction is considered a central feature of AD pathology that plays a significant role in disease progression, potentially independent of the Aβ-induced toxic effects.

The AD^2^ theory suggests the emerging role of Indoleamine 2,3-dioxygenase (IDO) in AD progression. IDO, an enzyme involved in tryptophan metabolism, plays a dichotomic role in neuroinflammation. It can either promote immune tolerance and anti-inflammatory responses or contribute to neurotoxic inflammation (11-13). In AD, elevated IDO activity may indicate a maladaptive immune response, amplifying neuroinflammation and accelerating neurodegeneration (14-15). This suggests that IDO could be a central mediator of the immune dysregulation seen in AD, linking the immune system’s response to the neuroinflammatory processes driving the disease.

Cytokines, particularly IFN-γ, and cGAS (cyclic GMP-AMP synthase) are key immune players that drive neuroinflammation in AD (14, 16-17). IFN-γ induces IDO expression, fostering a pro-inflammatory brain environment and activating microglia, which release additional cytokines and perpetuate chronic neuroinflammation (15-19). The cGAS-STING pathway, involved in immune responses to DNA damage, is also implicated in AD, where its activation in microglia triggers type I interferon responses, further intensifying inflammation and neurodegeneration (16,19). Together, IDO and cGAS form a synergistic feedback loop, amplifying inflammatory signals and exacerbating disease progression.

Thus, targeting IDO activity, along with cGAS modulation, could offer a novel approach for restoring immune balance in AD. By addressing both the immune dysregulation and neuroinflammatory components of the disease, therapies aimed at these pathways could provide a promising therapeutic strategy within the autoimmune framework of AD^2^, potentially offering disease-modifying effects beyond amyloid and tau-focused treatments.

Several studies have demonstrated that cannabidiol (CBD) can alleviate AD symptoms and improve patient outcomes (20-22). CBD, a non-psychoactive phytocannabinoid derived from cannabis, has gained attention as a promising therapeutic agent for inflammatory diseases, largely due to its anti-inflammatory properties, low toxicity, and ability to modulate various pro-inflammatory signals (22-23). Our findings suggest that CBD significantly increased TREM2 expression in glial cells and reduced IL-6 levels in peripheral blood leukocytes, which were associated with improved cognitive function in a pre-clinical AD model (24). Additionally, CBD not only enhanced astrocytic IL-33 expression, a cytokine that promotes microglial phagocytosis of amyloid-beta and improves contextual memory, but also boosted acetylcholine production, further improving cognitive performance (24). These results support the potential of CBD as a clinically safe and effective disease-modifying treatment to slow neurocognitive decline in AD.

CBD’s anti-inflammatory and immunomodulatory properties may influence key immune pathways such as IDO and cGAS, which are involved in chronic inflammation and neurodegeneration. While direct evidence of CBD’s interaction with these pathways is still limited, it is plausible that CBD could modulate both the IDO and cGAS pathways, offering potential therapeutic benefits for neuroinflammation in neurodegenerative diseases like AD. In this study, we aimed to investigate the association between CBD, IDO, and cGAS to uncover new insights into CBD’s role in modulating immune responses and slowing neurodegeneration, ultimately contributing to more effective, disease-modifying treatments for AD.

## Materials and methods

### Declaration Regarding Humane Use of Animals

All mouse experiments were conducted in accordance with the principles of the Three Rs’ (Replacement, Reduction, and Refinement) and were approved by the Institutional Animal Care and Use Committee (IACUC) at Augusta University (Augusta, GA, USA) under protocol #2011-0062. Procedures were designed to minimize discomfort and distress to the animals, and all animal care practices adhered to the highest standards of humane treatment as outlined by the NIH (National Institute of Health, Bethesda, Maryland, USA) guidelines.

### Experimental Design and Treatment Protocol

Adult male 5xFAD mice were purchased from Jackson Laboratory (Bar Harbor, Maine, USA). 5xFAD mice express human amyloid precursor protein (APP) and preseniln-1 (PSEN1) transgenes with five AD-linked mutations, used as a pre-clinical model of AD (25-26). At 9-12 months, mice were randomized into two groups (n = 10/group), receiving either placebo or inhaled CBD (10 mg/mouse, Thriftmaster Global Bioscience, Dallas, TX, USA) every day for 4 weeks. Each CBD inhaler contained 985 mg of broad-spectrum CBD (winterized crude hemp extract) plus 15 mg of co-solvent, surfactant, and propellant, with a total volume of 1000 mg (1.78 mg dose per actuation, with 200 mL/min flow rate). For the placebo, the 985 mg of broad-spectrum CBD was replaced with 985 mg of hemp seed oil. As described previously (27), inhalers were modified by adding an extra nozzle piece to adjust to the mouse model and to further control the intake of CBD. The study design included three independent cohorts, ensuring the robustness and reproducibility of the results.

### Immunofluorescence staining (IF)

Fresh brain tissues were fixed with 10% neutral buffered formalin. The tissues then were processed, embedded and subsequently cut into 4 μm sections for Immunofluorescence assessments as described previously (24). Briefly, after blocking endogenous peroxidase activity with hydrogen peroxide (diluted 1:10 in distilled water for 10 min), sections were treated with Proteinase K for 10 min and washed twice in PBS. Slides were then labeled with specific, fluorescent-conjugated antibodies with fluorescent conjugation against IDO (PE anti-IDO1, Biolegend USA, Cat# 654006), Transmembrane protein 119 (Alexa Fluor™ 488 anti TMEM119, for microglia, Thermo Fisher Scientific USA, Cat# 53-6119-82), Glial fibrillary acidic protein (Fluor™ 488 anti GFAP, for astrocytes, Biolegend USA, Cat# 644704) and cGAS (Cat#79978, purified, Cell signaling Technology, USA). All slides were counterstained using DAPI (4′,6-diamidino-2-phenylindole, Thermo Fisher Scientific USA, Cat# D1306) prior to examination and imaging by Zeiss Fluorescence Microscope. The integrated density of the area chosen (in pixels) and the mean gray value (the measurement of the brightness) were measured using lasso tool of Adobe Photoshop CS4 extended.

### Analytical flow cytometry

For flow cytometry analysis, single-cell suspension was prepared from brain by sieving the brain tissues through a 100μM cell strainer (BD Biosciences, San Diego, CA, USA) followed by centrifugation (252 g, 10 min). All cells were then stained with fluorescent antibodies based on routine flow cytometry staining protocol as described previously (24). Briefly, all cells were then stained with fluorescent antibodies to phenotype and quantify microglia (TMEM119^+^CD45^+/lo^) and infiltrating macrophages (TMEM119^-^CD45^+/hi^) (Alexa Fluor™ 488 anti TMEM119, Thermo Fisher Scientific USA, Cat# 53-6119-82 & PE anti-mouse CD45, Biolegend USA, Cat# 157604). Then cells were fixed and permeabilized and stained intracellularly for cytokines including Interferon gamma (APC anti-mouse IFNγ, Biolegend USA, Cat#505809), Interleukin-1β (PerCP anti-mouse IL-1β, Thermo Fisher Scientific USA, Cat# 46-7114-82), Tumor Necrosis Factor alpha (Pacific Blue™ anti-mouse TNFα, Biolegend USA, Cat#506318) and Interleukin-10 (Biotin anti-mouse IL-10, Biolegend USA, Cat#505003). Cells were then run through a NovoCyte Quanteun flow cytometer (Agilent Technologies, Santa Clara, CA, USA) and analyzed by FlowJo V10 analytical software. To confirm the specificity of primary antibody binding and rule out nonspecific Fc receptor binding to cells or other cellular protein interactions, negative control experiments were conducted using isotype controls matched to each primary antibody’s host species, isotype, and conjugation format.

### Integrated bioinformatic analysis

#### Network construction: protein–protein interaction

To investigate the IDO/cGAS genes, we used the interactive database platform STRING v.11.0 (https://string-db.org/). STRING is a database of known and predicted protein-protein interactions (28). The interactions include direct (physical) and indirect (functional) associations; they stem from computational prediction, from knowledge transfer between organisms, and from interactions aggregated from other (primary) databases. Next, the protein–protein inter-action (PPI) network was constructed. The confidence score cutoff was set at 0.4, and other settings were set to default.

#### Gene ontology functional analysis

After identification of the IDO/cGAS targets, the Metascape platform and the Enrichr database were used to analyze their main biological processes, cellular components and molecular functions to perform enrichment analysis (28-29). The results were visualized using biological online tools.

### Statistical analysis

For statistical analysis, Brown-Forsythe and Welch analysis of variance (ANOVA) was used to establish significance (p < 0.05) among groups. For tissue quantification statistical analysis, the area of expression was compared in both placebo and CBD treated groups using two-way ANOVA, followed by post-hoc Sidak testing for multiple comparison (p < 0.05).

## Results

### Inhaled CBD Reduces IDO Expression and Alters glial Activation in mice with AD

Immunofluorescence staining was performed to evaluate the effects of inhaled CBD on IDO expression in the brains of 5XFAD mice, a well-established model of AD. A significant reduction in IDO expression was observed in entorhinal cortex area of both microglia and astrocytes in CBD-treated mice compared to untreated controls (Figure 1). Specifically, in panel A, TMEM119+ microglial cells showed a notable decrease in IDO expression following CBD treatment, while panel B demonstrated a similar reduction in IDO levels in GFAP+ astrocytes. Quantitative analysis, shown in panel C, further confirmed these observations, with pixel intensity measurements revealing a significant decrease in total IDO expression as well as in both microglia and astrocytes in CBD-treated AD mice (***p < 0.001). In addition to these reductions in IDO expression, pixel intensity analysis indicated changes in glial activation. Specifically, CBD treatment was associated with increased microglial activation, as reflected by a higher pixel intensity of TMEM119+ staining, suggesting a heightened state of microglial response. In contrast, astrocyte activation, measured by GFAP intensity, was reduced in CBD-treated animals compared to controls, indicating a potential modulatory effect of CBD on astrocyte function. While the differences in both microglial and astrocyte activation were non-significant (ns), the observed trends, higher microglial activation and lower astrocyte activation, are significant in the context of AD, as increased microglial activation and reduced astrocyte activation are considered beneficial for AD progression. These changes support the potential therapeutic effects of CBD in modulating neuroinflammatory pathways in AD.

**Fig 1.**
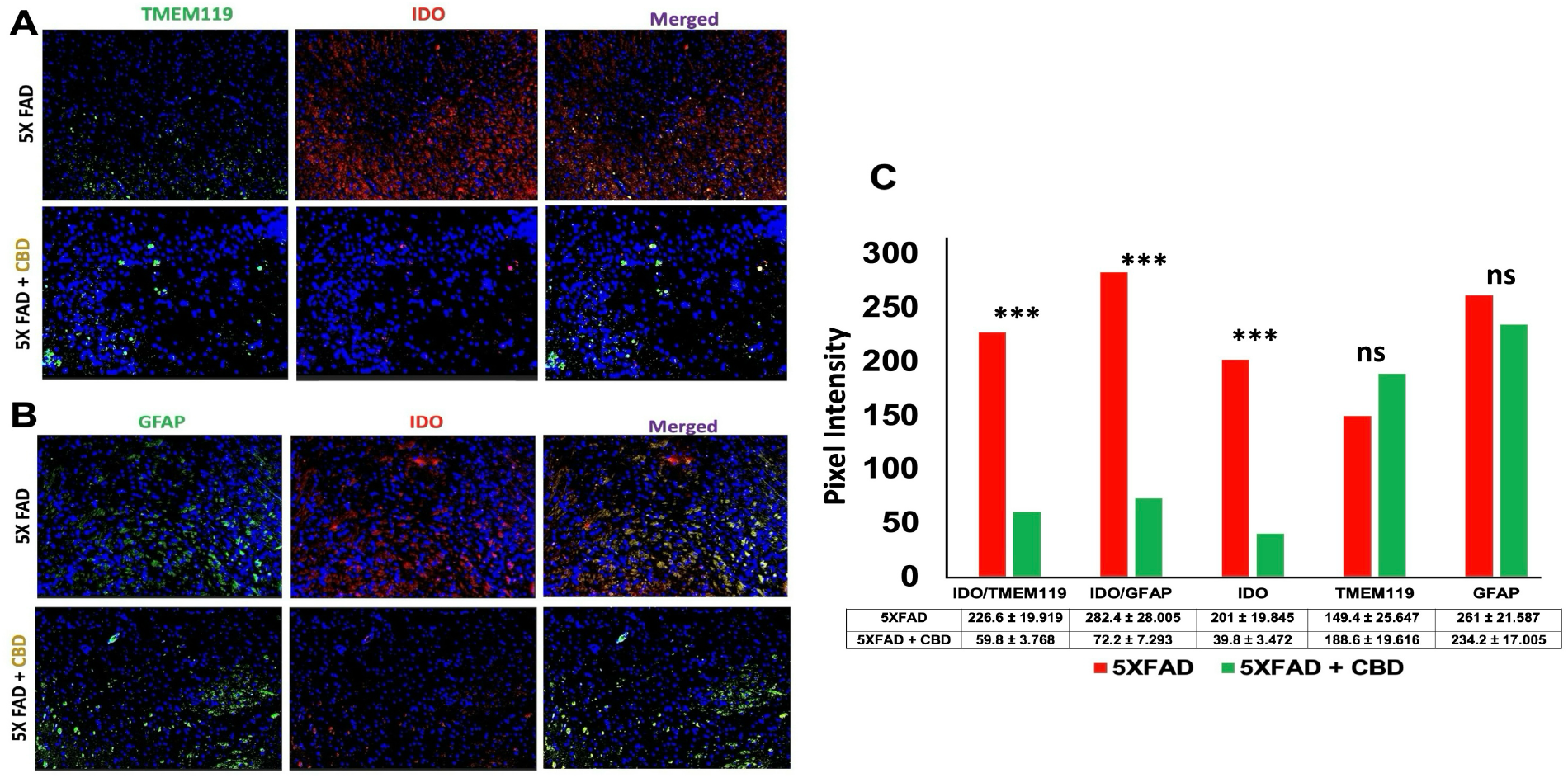
Inhalation of CBD Reduces IDO Expression in 5xFAD Mice with AD. Immunofluorescence staining (panel **A** and **B**) demonstrated a significant reduction in IDO expression in microglia (TMEM119+ cells, panel A) and astrocytes (GFAP+ cells, panel B) within the entorhinal cortex of CBD-treated 5xFAD mice, compared to untreated controls. The images are representative of three independent cohorts of experiments. **C)** Quantitative analysis of IDO expression, based on pixel intensity measurements, confirmed a significant decrease in total IDO expression as well as in both microglia and astrocytes in CBD-treated mice with Alzheimer’s disease (***p < 0.001). Additionally, pixel intensity analysis revealed increased microglial activation (higher TMEM119 intensity) and reduced astrocyte activation (lower GFAP intensity) in CBD-treated animals, relative to untreated controls, though differences in microglial and astrocyte activation were non-significant (ns).

The reduction in IDO expression, particularly in microglia and astrocytes, suggests that CBD may modulate key inflammatory pathways involved in AD pathology. IDO, a crucial enzyme in tryptophan metabolism, is known to be upregulated in neuroinflammatory conditions and has been implicated in neurodegenerative diseases like AD. By reducing IDO expression, CBD could potentially mitigate neuroinflammation and its contribution to AD progression, highlighting its therapeutic potential in targeting the immunoinflammatory aspects of the disease.

### CBD Modulates the IDO/cGAS Pathway in mice with AD

Immunofluorescence staining revealed that inhalation of CBD significantly decreased cGAS expression in IDO-expressing cells in entorhinal cortex of 5XFAD mice with AD, compared to untreated controls (Fig. 2A). This reduction in cGAS expression was quantified by measuring the co-localization of cGAS and IDO, with pixel intensity analysis showing a marked decrease in cGAS/IDO co-expression in CBD-treated AD mice relative to untreated controls (Fig. 2B; p < 0.0001). These findings suggest that CBD may offer a promising therapeutic approach for AD by targeting neuroinflammatory mechanisms, specifically the IDO/cGAS pathway. This result has two key implications: First, it supports the autoimmune theory of AD (AD^2^), providing evidence that a more immune-targeted treatment strategy could emerge for AD, moving beyond the traditional focus on amyloid-beta. Second, our work introduces a novel mechanism through the modulation of the IDO/cGAS pathway, which warrants further investigation into CBD’s potential to alter neuroinflammation and immune responses in AD. These results underline the importance of CBD as a candidate for treating AD via immune-based pathways, expanding our understanding of its therapeutic potential in neurodegenerative diseases.

**Fig 2.**
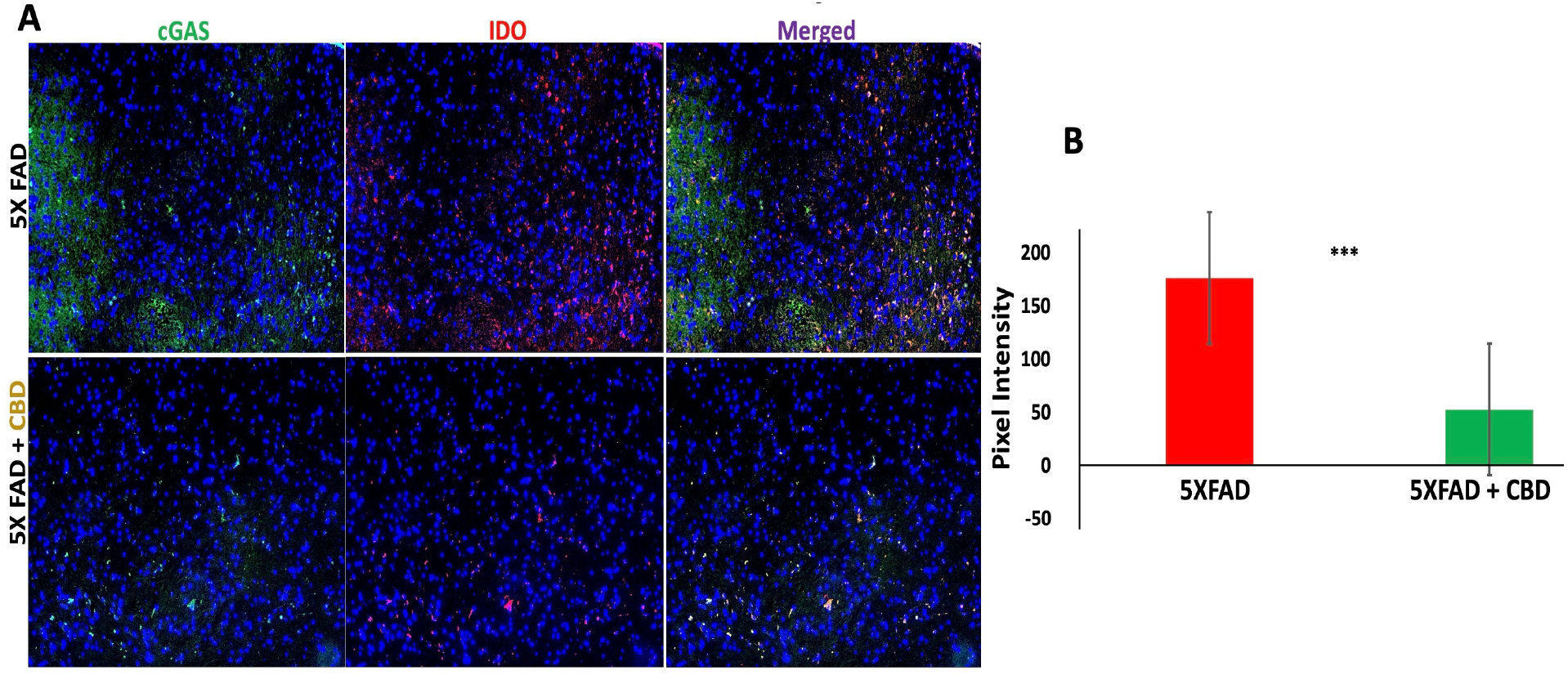
Inhalation of CBD Reduces cGAS/IDO Co-Expression in 5xFAD Mice with AD. **A)** Immunofluorescence staining showed that CBD inhalation significantly reduced cGAS expression in IDO-expressing cells within the entorhinal cortex of 5xFAD mice with Alzheimer’s disease, compared to untreated controls. The images are representative of three independent cohorts of experiments. **B)** Quantitative analysis of co-localization between cGAS and IDO, based on pixel intensity measurements, revealed a significant decrease in cGAS/IDO co-expression in CBD-treated mice with Alzheimer’s disease, relative to untreated controls (***p < 0.0001).

### Inhaled CBD Regulates Proinflammatory Immune Profile in Mice with AD

Flow cytometry analysis was performed to assess immune cell profiles in the brains of 5XFAD mice with or without CBD treatment (Figure 3). Panel A shows the total brain cells from untreated (upper panel) and CBD-treated (lower panel) 5XFAD mice, with live gating based on FSC/SSC parameters. Panel B reveals that CBD treatment significantly reduced the frequency of infiltrating macrophages (CD45+/hiCD11b+TMEM119-) compared to untreated controls, indicating a decrease in neuroinflammatory responses. Further analysis (Panel C) showed that CBD treatment decreased proinflammatory cytokine production, while enhancing the anti-inflammatory cytokine IL-10. Panel D quantifies these changes, with significant differences (**p<0.05, ***p<0.001) observed in the cytokine profile. These results suggest that CBD modulates immune responses in AD by reducing macrophage infiltration and promoting an anti-inflammatory cytokine profile, supporting its potential as a therapeutic agent for neuroinflammation in AD.

**Fig 3.**
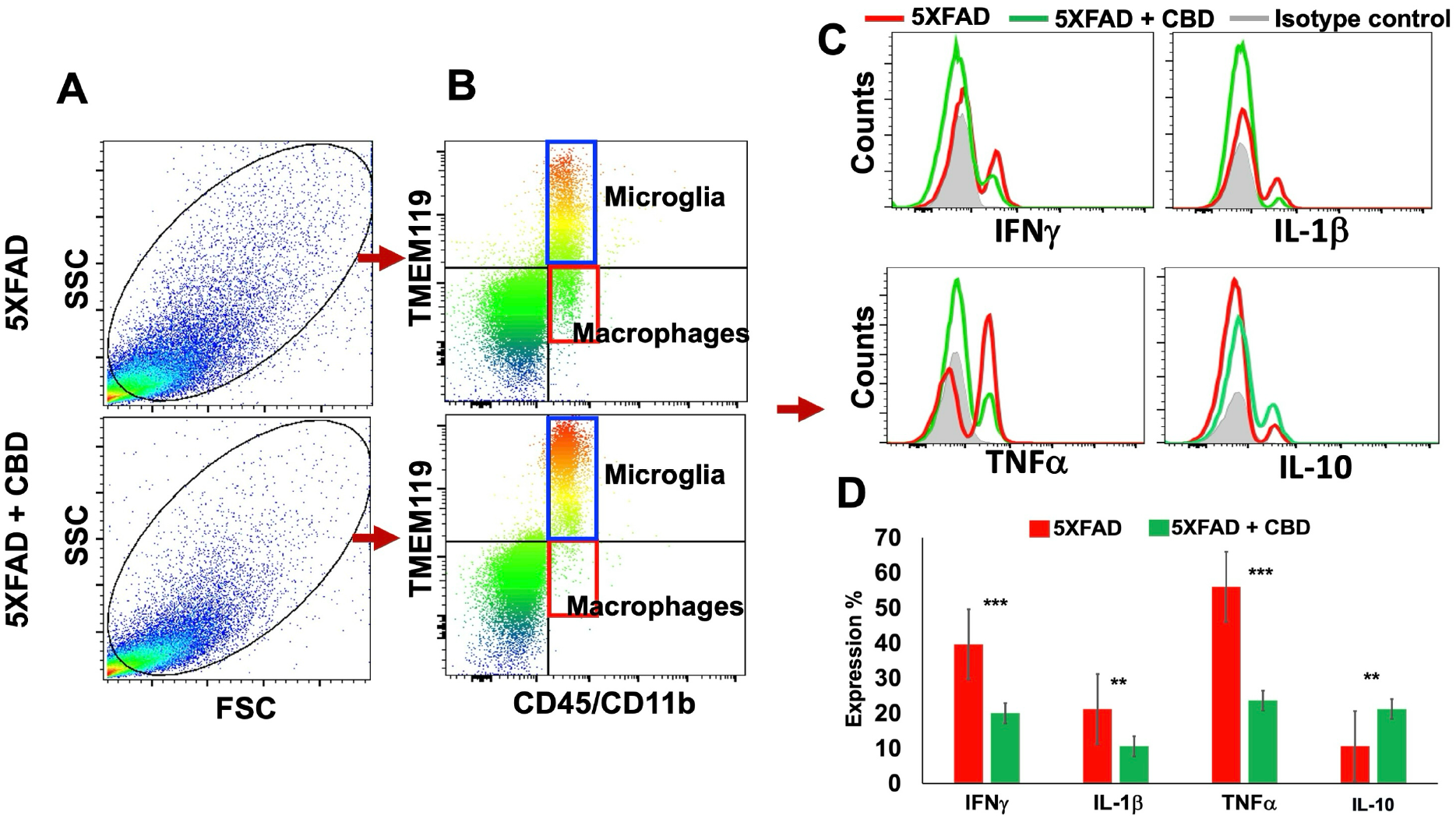
Inhaled CBD Modulates the Proinflammatory Immune Profile in Mice with AD. **A)** Flow cytometry dot plots show total brain cells from untreated (upper panel) and CBD-treated (lower panel) 5xFAD mice, with live gating based on forward scatter (FSC) and side scatter (SSC) parameters. Data represent three independent cohorts of experiments. **B)** Flow cytometry analysis distinguishes resident microglia (CD45^+/lo^CD11b^+^TMEM119^+^) from infiltrating macrophages (CD45^+/hi^CD11b^+^TMEM119^-^). CBD treatment reduced the frequency of infiltrating macrophages, indicating a decrease in inflammatory responses. **C)** Flow cytometry analysis reveals a significant reduction in proinflammatory cytokine production in CBD-treated mice, along with enhanced production of the anti-inflammatory cytokine IL-10 compared to untreated controls. **D)** Cytokine profile alterations were quantified and visualized in histograms, with significant differences observed (**p<0.05, ***p<0.001).

### Bioinformatics Analysis of IDO/cGAS Pathway in AD: CBD as a Potential Therapeutic Target

To explore the potential therapeutic role of CBD in Alzheimer’s disease (AD), we conducted bioinformatics analysis focusing on the IDO/cGAS pathway and its interaction with CBD (Figure 4). Panel A presents a protein-protein interaction (PPI) network analysis, which identifies key genes, such as IDO and cGAS, as central components in neuroinflammatory pathways associated with AD. Additionally, the analysis highlights several potential CBD-targeted molecules, including AKT1, TRPV1, and GPR55, which may play critical roles in modulating these pathways. Panel B further explores the PPI enrichment analysis, which reveals co-regulation between the IDO/cGAS pathways and CBD’s molecular targets. This co-regulation suggests that CBD may exert its therapeutic effects by influencing interconnected molecular networks that are essential for cellular function and immune modulation. Panel C illustrates the results of pathway enrichment analysis, which shows that CBD targets genes involved in critical biological processes, such as the regulation of nitric oxide synthase, immune system modulation, and calcium ion concentration. These processes are pivotal for maintaining neuroinflammatory balance and preventing AD-associated neurodegeneration. Taken together, these bioinformatics findings suggest that CBD holds promise as a therapeutic agent targeting the IDO/cGAS pathway and associated molecular pathways, potentially offering new approaches for modulating neuroinflammation and mitigating AD progression.

**Fig 4.**
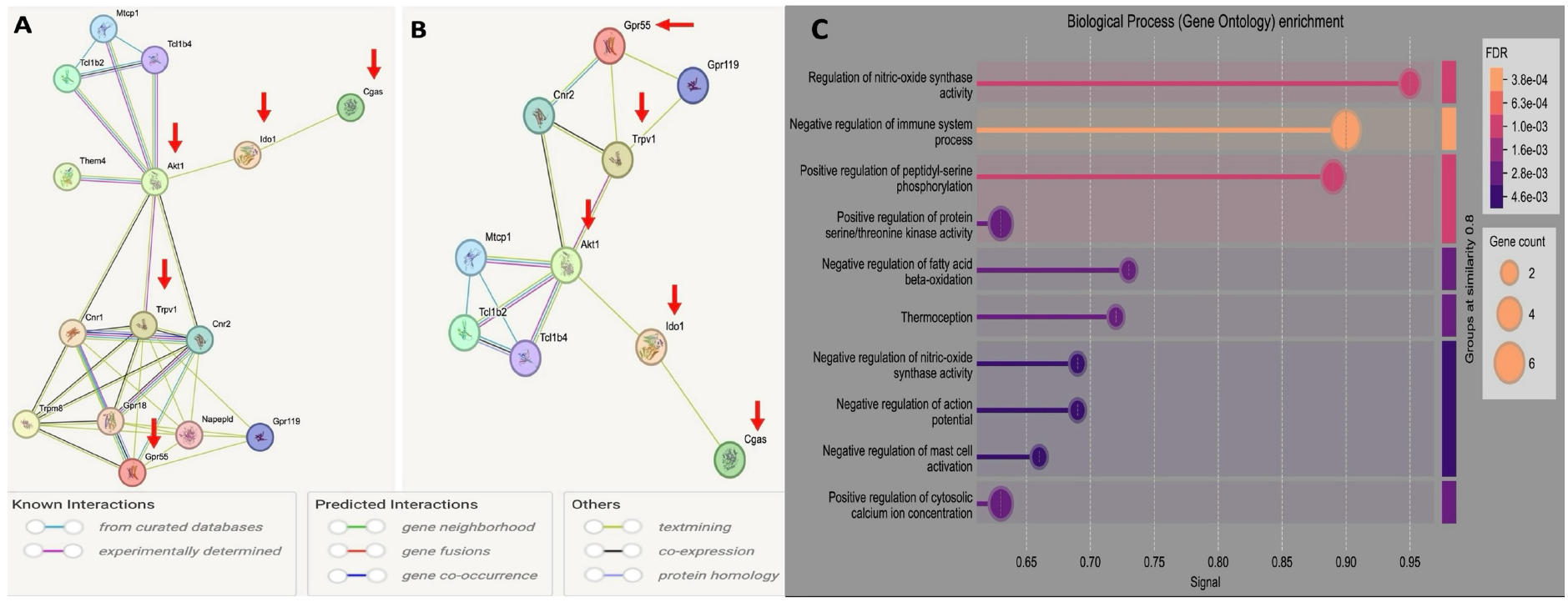
Bioinformatics analysis of IDO/cGAS Pathway in AD: CBD as a potential Therapeutic Target. **A)** Protein-protein interaction (PPI) network analysis reveals key genes, IDO and cGAS, along with potential CBD-targeted molecules (AKT1, TRPV1, GPR55). **B)** PPI enrichment analysis suggests co-regulation between IDO/cGAS pathways and CBD’s molecular targets, highlighting their involvement in crucial cellular processes. **C)** Pathway enrichment analysis indicates that CBD targets genes involved in biological processes such as regulation of nitric oxide synthase, immune system modulation, and calcium ion concentration, all of which are critical for maintaining neuroinflammatory balance and preventing AD-related inflammation and neurodegeneration.

## Discussion

Our study provides robust support for the evolving concept of Alzheimer’s disease as an autoimmune disorder (AD^2^), offering new insights into the pathophysiology of AD beyond the amyloid-β hypothesis (7,30-31). While amyloid plaques and tau tangles have historically been considered central to AD pathology, emerging evidence suggests that these factors alone cannot fully account for the complexity of AD progression (32-34). Our findings are consistent with the immune hypothesis of AD^2^, which proposes that immune system dysregulation, rather than amyloid deposition in isolation, plays a central role in the initiation and progression of neurodegenerative processes in AD (7,30-34). Specifically, in this model, the immune system aberrantly targets autologous brain tissue, exacerbating neuroinflammation, a key feature of AD pathology (7,35-36). By demonstrating that CBD inhalation modulates immune-related pathways in the brain, our results provide compelling evidence supporting immune dysfunction as a major contributor to AD progression, further validating the AD^2^ model.

The choice of the 5XFAD transgenic mouse model for our studies was particularly advantageous, as it closely mimics the hallmark features of AD, including amyloid-β deposition, neuroinflammation, and cognitive decline (25-26). This model is widely recognized for its ability to reproduce key aspects of AD pathology, making it an ideal system for evaluating potential therapeutic strategies like CBD. The 5XFAD model provided a robust platform to investigate how CBD interacts with immune-related pathways in the context of AD, allowing us to generate meaningful insights into its effects on neuroinflammation and glial function.

A critical mechanism implicated in immune dysfunction within the CNS is the IDO/cGAS pathway (16,37-38). Both IDO and cGAS are integral to maintaining immune homeostasis in the brain (11-13,16). IDO regulates tryptophan metabolism, and its overactivation is associated with increased neuroinflammation in various neurological conditions, including AD (39-40). cGAS, through its role in sensing cytosolic DNA, initiates the STING pathway, which, when dysregulated, can amplify neuroinflammation (41-42). Our study presents novel evidence that CBD effectively modulates key components of immune regulation, highlighting its potential as a therapeutic strategy to address immune dysfunction at the core of AD pathophysiology. By targeting both IDO and cGAS, CBD offers a promising strategy to restore immune balance in the CNS and mitigate neurodegenerative processes.

Our findings are in line with previous studies that have shown beneficial effects of CBD in AD models, both in mice and humans (20-24). These studies have demonstrated that CBD’s anti-inflammatory and neuroprotective properties can ameliorate key aspects of AD pathology, such as amyloid plaque deposition and cognitive impairment. Our work builds on this foundation, specifically highlighting the modulation of immune pathways like IDO/cGAS as a novel mechanism by which CBD could exert its therapeutic effects in AD. CBD’s modulation of the IDO/cGAS pathway is accompanied by significant effects on glial cell activity. Our results demonstrate that CBD treatment reduces the expression of IDO in key glial populations, including microglia and astrocytes, while also decreasing the co-expression of IDO and cGAS. Given the central role of glial cells in modulating immune responses within the CNS (43), these findings suggest that CBD directly influences glial-mediated immune regulation. In conjunction with this, we also observed a CBD-induced increase in microglial activation alongside a reduction in astrocytic activation. This dual modulation is significant, as elevated microglial activation and lower astrocytic activity have been associated with enhanced neuroprotection and improved outcomes in AD (44). Chronic neuroinflammation, driven by the overactivation of immune pathways like IDO and cGAS, plays a pivotal role in AD progression (39-42). Therefore, CBD’s ability to regulate both glial activity and immune pathways may not only mitigate neuroinflammation but also foster a neuroprotective environment within the brain, offering a potential therapeutic benefit for AD.

Our bioinformatic analysis provides further insight into the mechanisms underlying CBD’s therapeutic potential in AD. Notably, the analysis suggests that CBD may promote neuronal survival through activation of the AKT1 signaling pathway, a critical pathway involved in cell survival, metabolism, and neuroprotection (45). In addition, CBD appears to modulate key receptors such as TRPV1 and GPR55, which are implicated in neuroinflammation, pain management, and neuroprotection (46). This multifaceted action further supports the idea that CBD is not only addressing neuroinflammation but also promoting neuronal resilience, thereby enhancing its therapeutic potential in AD.

The ability of CBD to modulate both immune dysfunction and neuroinflammation offers a more comprehensive approach to AD treatment compared to traditional amyloid-based therapies, which have shown limited success in clinical trials (47). CBD’s effects on the IDO/cGAS pathway, coupled with its ability to influence glial activity and neuronal survival, position it as a promising candidate for AD therapy. Moreover, CBD’s impact on cytokine profiles further underscores its therapeutic potential. In our study, CBD treatment was associated with a decrease in pro-inflammatory cytokines, such as IFN-γ, IL-1β, and TNF-α, while increasing the production of the anti-inflammatory cytokine IL-10. This modulation of the inflammatory milieu is particularly important in AD, where chronic neuroinflammation plays a central role in disease progression (48-50). By restoring immune homeostasis and attenuating neuroinflammation, CBD may help slow or halt disease progression, providing a valuable addition to current therapeutic strategies.

In conclusion, our findings contribute to the growing body of evidence supporting the immune theory of AD^2^, demonstrating that CBD modulates key immune pathways, particularly the IDO/cGAS pathway, which plays a critical role in immune dysregulation in AD. Our results highlight the potential of CBD as a novel therapeutic agent for AD, offering a multifaceted approach to disease management. By modulating immune responses, regulating glial activity, and promoting neuronal survival, CBD addresses several key aspects of AD pathology. These findings warrant further clinical investigation into the use of CBD, either as a monotherapy or in combination with other therapeutic strategies, to evaluate its potential for modifying the course of AD.

### Limitations

Two major limitations of this study should be noted. First, we used only one dose of CBD in the 5XFAD mice model. We acknowledge that testing multiple doses could strengthen the conclusions and provide a more comprehensive understanding of CBD’s therapeutic potential. Dose optimization is an important aspect of CBD-based therapies, and future studies should explore a range of doses to determine the most effective treatment regimen. Second, while this study offers valuable preclinical data, the translation of cannabidiol doses from mice to humans remains a challenge due to species differences in metabolism and pharmacokinetics. We have addressed the human equivalence issue in the limitations section to highlight the need for clinical trials to determine the optimal CBD dose for AD treatment in humans. These concerns emphasize the importance of further research and human trials to confirm the therapeutic potential of CBD.

In addition, while we focused on macrophages and microglia as key inflammatory cell types, evaluating a broader range of cellular responses to CBD, including the role of astrocytes and other immune cells, would provide a more comprehensive understanding of its effects on neuroinflammation. Future research should also explore how CBD may influence other aspects of neuroinflammation, particularly by examining the broader cellular landscape beyond macrophages and microglia. These areas represent important directions for future studies, and we plan to explore them in more depth in subsequent research.

## Statements & Declarations

## Acknowledgements

Authors are thankful to Thriftmaster Holding Group for providing the inhalant CBD for this study. Authors also thank Medicinal Cannabis of Georgia for providing help in the analysis, processing the data and optimizing the CBD dosage.

## Author contributions

LPW, ELS and BB: Study conception and design, analysis and interpretation of data, and drafting the article., NPM, ELS, SEN, LMM, BH, HK, and BBH: Acquisition of data and drafting the article., HMR, NY, JCY, DCH, and KMD: Editing and Scientific Contribution.,

## Funding

This work was supported by institutional seed funding from the Dental College of Georgia at Augusta University.

## Competing interests

(1) Lei Phillip Wang, Babak Baban, and Jack Yu are members of Medicinal Cannabis of Georgia with no financial interest. (2) All other authors declare no conflict of interest. (3)Thriftmaster Holding Group (THG) is the provider of CBD inhalers and has a licensing contract with Augusta University. (4) THG had no role in study design, data collection and analysis, decision to publish, or preparation of the manuscript.

## Data availability

The datasets generated during and/or analyzed during the current study are available from the corresponding author on reasonable request.

## Ethics approval

Animal experiments were approved by the Institutional Animal Care and Use Committee (IACUC) of Augusta University and followed the IACUC guidelines. There was no human subjects/samples used in this study.

